# A unique neural signature of long-term memory encoding from EEG inter-electrode correlation

**DOI:** 10.1101/2025.10.21.683799

**Authors:** Chong Zhao, Edward K. Vogel, Monica D. Rosenberg

## Abstract

Classic memory models proposed that the encoding process involved in visual working memory (VWM) controls the bandwidth of encoding in long-term memory (LTM). Behaviorally, VWM and LTM accuracies are reliably correlated at the behavioral level, raising the question of whether LTM encoding uniquely engages processes that are distinct from VWM encoding. To investigate this, we recorded EEG activity as participants completed recognition memory tasks with set sizes of 32 and 128, far beyond typical VWM capacity. Using interelectrode correlation (IC) analysis, we found that IC patterns reliably predicted individual differences in LTM encoding across both set sizes, indicating a robust, domain-general neural signature. Importantly, this predictive power remained even after controlling for VWM and attentional control performance, suggesting that the model captures variance specific to LTM encoding. Temporally, predictive signals emerged only after stimulus onset and persisted for 500–600 ms. Early and late encoding phases involved distinct network structures, reflecting dynamic neural processes underlying individual differences in LTM encoding. Together, our findings reveal a unique and temporally dynamic neural signature that supports individual differences in LTM encoding, independent of general cognitive abilities.

**Significance Statement:** Humans maintain visual information using two distinct memory systems: visual working memory, which is capacity-limited, and long-term memory, which has virtually unlimited capacity. Despite their differences, individuals with better VWM performance often show superior LTM performance, raising the question of whether unique LTM encoding abilities exist beyond those accounted for by VWM. Here, we leveraged EEG data to examine interelectrode correlation patterns during memory encoding. Our results show that these neural patterns uniquely predict individual differences in LTM encoding ability, even after controlling for VWM performance. Furthermore, we showed that the interelectrode correlation patterns continuously track the temporal dynamics of LTM encoding over time.

## Introduction

Imagine sitting in a movie theater: the film currently playing occupies your conscious awareness, while memories of a previous visit to the same theater may also be brought back into mind. To distinguish these two types of experiences, William James (1890) introduced the concepts of primary memory and secondary memory. Primary memory refers to information actively held in consciousness, whereas secondary memory involves information that must be retrieved into consciousness for use. Building on this distinction, the classic modal model of memory (Atkinson & Shiffrin, 1968) proposed that information first enters a short-term store (i.e., primary memory) and, through processes such as rehearsal, may subsequently be encoded into a long-term store (i.e., secondary memory).

To capture how short-term store is used in real life, the term working memory is now used to describe the capacity-limited system that supports the maintenance and manipulation of information on short time scales. Notably, this capacity has been estimated at approximately four items (Cowan, 2001), a limit that is reflected in neural markers of visual working memory (VWM) load, which tend to plateau around this number (Vogel & Machizawa, 2004). When the number of items exceeds working memory capacity, it is assumed that excess items may be offloaded into visual long-term memory (LTM), which is thought to have virtually unlimited capacity. Indeed, empirical studies have demonstrated that individuals can store thousands of unique objects in visual LTM (e.g., over 2,000 items for 30 minutes in Brady et al., 2008; up to 10,000 scenes for ∼10 hours in Lionel, 1973), and do so with remarkable spatiotemporal precision (Wolfe et al., 2023).

Despite the stark contrast in capacity limits between working and long-term memory, individual differences in these systems are often moderately correlated. For example, Unsworth et al. (2014) found that although VWM and LTM performance loaded onto distinct latent factors, the two were moderately correlated across individuals, an effect that has been replicated with simple item memory as well as source memory judgments (Zhao & Vogel, 2025b, 2025a). One hypothesis for this relationship is that VWM and LTM share a common encoding bottleneck, such that limitations in initial encoding processes constrain both systems. Supporting this view, Fukuda and Vogel (2019) demonstrated that VWM capacity predicted LTM performance when participants were required to encode supracapacity arrays (e.g., 4 or 6 objects) in a VWM task, with LTM tested after a delay. Encoding bottlenecks in both memory stores may be affected by attentional control (AC) in that people with fewer lapses show better VWM capacity (Adam et al., 2015) and LTM task performance (Zhao et al., 2025).

Thus, VWM and LTM may share the same encoding constraints despite their vastly different capacity limits. This leaves an open question: is there a distinct neural signature of LTM encoding ability that predicts individual differences in LTM, independent of VWM abilities? A core criterion for identifying a neural marker of LTM encoding is that it should predict individual differences in LTM performance. Furthermore, if LTM encoding engages neural processes distinct from those underlying VWM, then such a marker should continue to predict LTM performance even after regressing out VWM and attentional control abilities. Conversely, if VWM and LTM share a common neural mechanism during encoding, regressing out VWM-related variance will significantly reduce the predictive power of the LTM encoding model.

In the current study, we recorded electroencephalogram (EEG) signals while 111 participants performed a visual LTM encoding task involving large set sizes (32 and 128 items), far exceeding typical VWM capacity. In a separate session after EEG, participants completed VWM and attentional control tasks. To capture the neural dynamics of encoding, we employed a recently developed EEG-based modeling approach, interelectrode correlations (Hakim et al., 2021), which quantifies large-scale correlational patterns across the scalp during task performance. We trained predictive models on the neural activities recorded during the encoding phase of the LTM task and used them to predict individual differences in subsequent LTM performance for left-out subjects. Specifically, we tested whether the trained LTM encoding model could (1) capture a common LTM encoding ability across different set sizes (32 and 128) and (2) uniquely predict LTM performance even after controlling for VWM capacity and attentional control, both measured independently in separate tasks.

## Method

### Participants

One hundred and eleven participants took part in the study (mean age = 25.1 years). Sample size was determined using G*Power, aiming to achieve 80% power to detect a small effect size (r = 0.3) at the conventional alpha level of .05. All participants reported normal or corrected-to-normal vision and normal color perception, and no history of neurological disorder. Study procedures were approved by the relevant University of Chicago Institutional Review Board and participants were compensated for their participation.

### Stimuli and Procedure

#### Day 1: Recognition Memory Task with EEG (set size 32 and 128)

Images were selected from the THINGS dataset (Hebart et al., 2023), a database with 1854 unique concepts and 26107 images. One exemplar was selected from each concept so that no repeated concept was used throughout each session (i.e., apple versus avocado). The recognition memory procedure followed established protocols in the literature and consisted of separate study and test phases (Fukuda & Woodman, 2015; Zhao & Woodman, 2020). In the set size 32 condition, participants viewed a stream of 32 images, each presented sequentially at the center of the screen for 800 ms. Stimuli were displayed against a gray background (90.0 cd/m²). A white fixation cross was superimposed on each image to help maintain central fixation and discourage eye movements. The interstimulus interval between images was randomly jittered between 250 and 400 ms to reduce anticipatory responses. Following the encoding phase, participants completed a series of recognition test trials. On each test trial, a single image was displayed at the center of the screen for 800 ms, again with a central white fixation cross. To minimize motor and ocular artifacts, participants were instructed to remain still and refrain from any movements while the fixation cross was visible. After 800 ms, the fixation cross disappeared, and the image remained on screen until a response was made. Half of the test images had been shown during the preceding study phase (old), while the other half were entirely new (new). The order of old and new images was randomized across trials. Participants were asked to indicate whether the image was “old” (previously studied) or “new” (not previously seen) using designated keyboard keys (“z” for old, “/” for new). In the set size 128 condition, participants learnt 128 images during the encoding phase, and 128 old and 128 new images were shown during the test phase. The trial structure and response key are the same as the set size 32 condition. Each participant completed 4 blocks of set size 32 and 1 block of set size 128, and the order of these blocks are randomized. In this EEG session, we also administered blocks with smaller set sizes (1, 2, 4 and 8), but since set size 32 and 128 are the ones that uniquely require LTM encoding processes, we did not include any of the smaller set sizes into our analyses. For a subset of our subjects that returned for a third online session, we found that LTM performance in the booth predicted LTM performance measured online (r(78) = 0.51, p < 0.01, offline measure set size 100, stimuli set from Brady et al., 2008).

**Fig 1.**
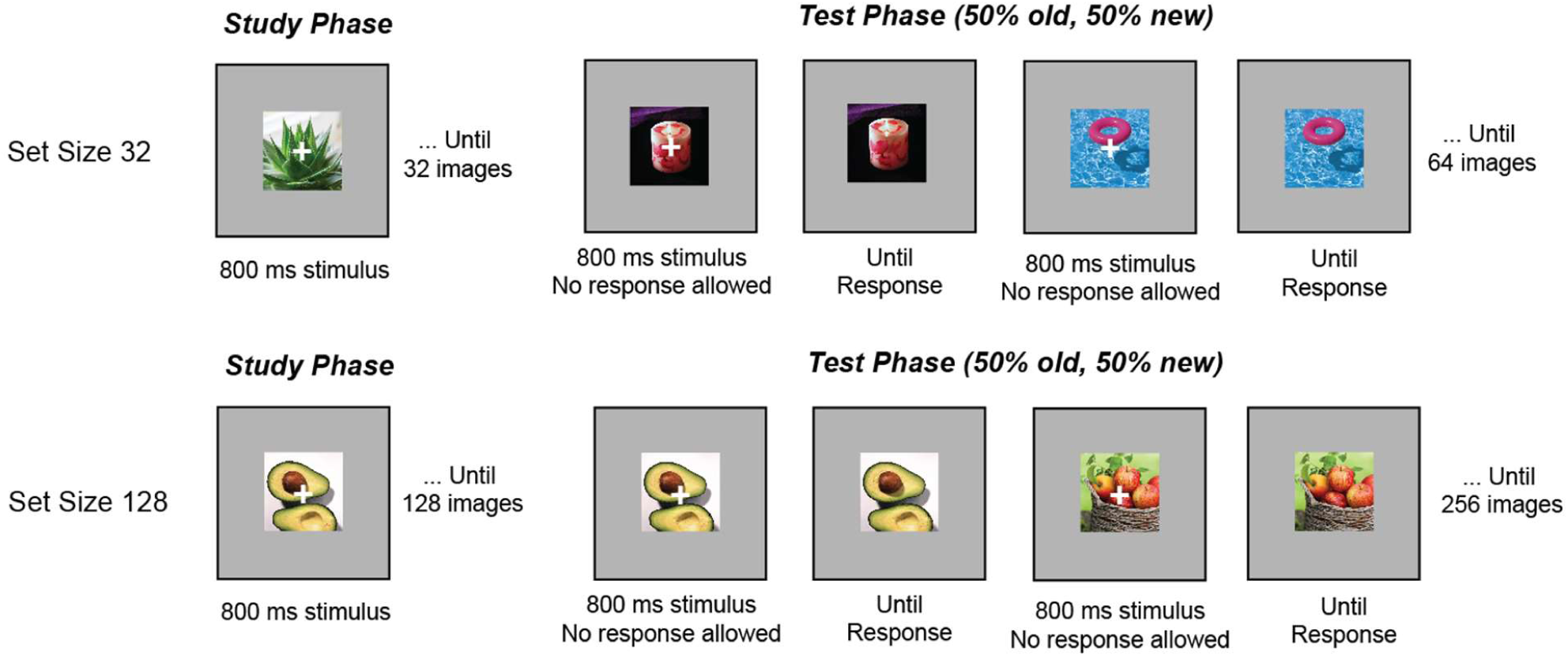
Visual Recognition Memory Paradigm with set size 32 and set size 128 with EEG.

#### Day 2: Online Attentional Control and Visual Working Memory tasks (no EEG)

After the EEG session, we emailed participants an invitation to complete an online battery of two attentional control tasks and two visual working memory tasks. Out of the one hundred and eleven participants, eighty participants consented to participate in this online study, with an average gap of 35.2 days.

##### Visual Working Memory Tasks

###### (1) Change Localization

The Change Localization task was derived from the color Change Localization task utilized in previous studies (Zhao et al., 2022, see Fig. 2A). In each trial, six colored squares were presented simultaneously for 250 ms, followed by a blank retention interval of 1,000 ms. Subsequently, the same six squares reappeared in their original positions, with one color altered to a hue not previously presented in that trial. Each square was numbered from 1 to 6, and participants were instructed to press the corresponding key to identify the square that had changed color. The spatial arrangement of the six numbers was randomized across trials. Our behavioral measure of interest was accuracy in the Change Localization task.

**Fig 2.**
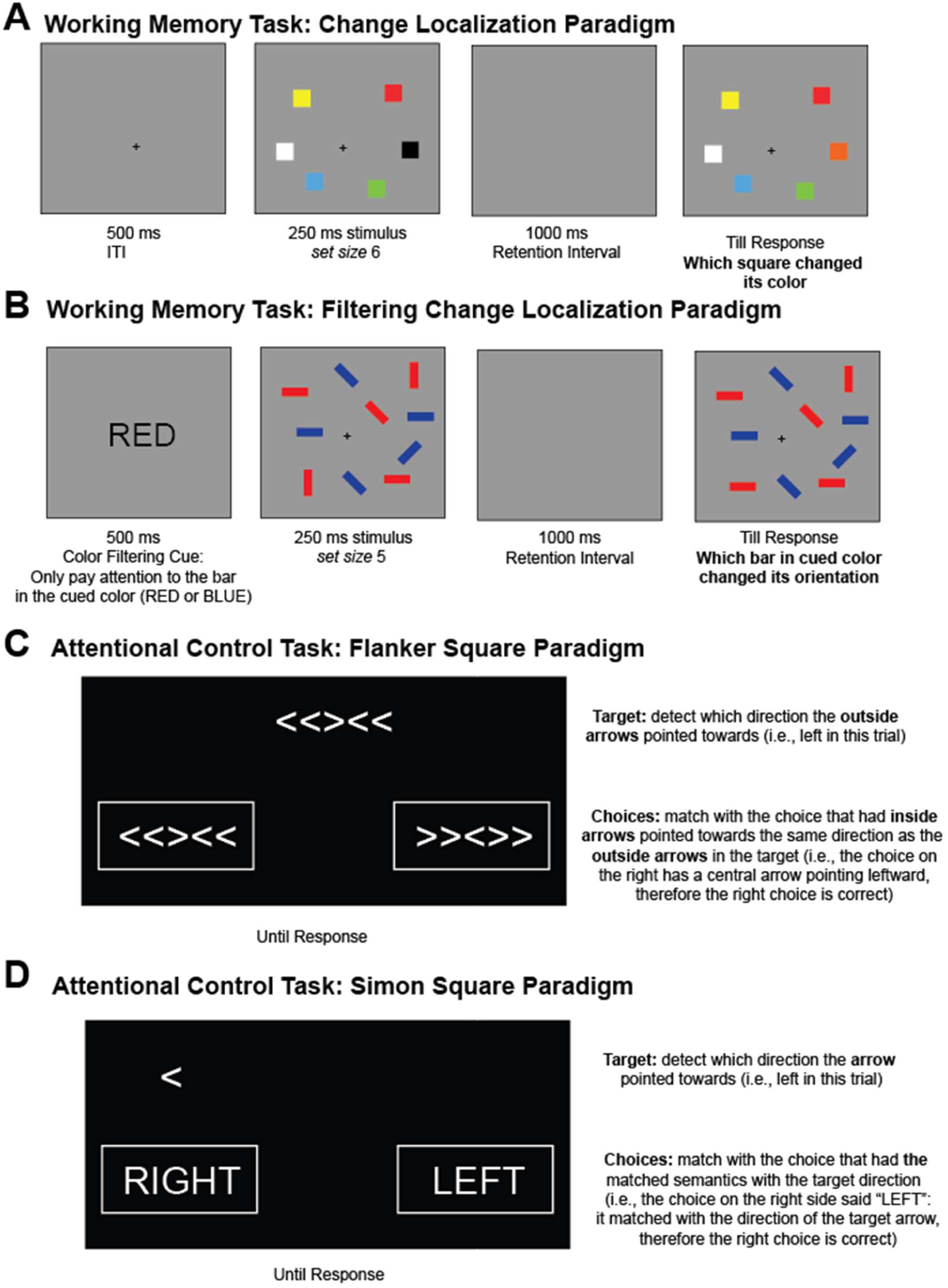
Online Working Memory and Attentional Control Tasks (Adapted from Zhao & Vogel, 2025). (A) Schema of Change Localization Task (Adapted from Zhao, Vogel & Awh, 2023). (B) Schema of the Filtering Change Localization Task (Adapted from Zhao & Vogel, 2025). (C) Schema of the Flanker Square Task (Adapted from Burgoyne et al., 2023). (D) Schema of the Simon Square Task (Adapted from Burgoyne et al., 2023).

###### (2) Filtering Change Localization

The Filtering Change Localization task was adapted from the Filtering Change Detection task (Luck & Vogel, 1997; Zhao & Vogel, 2025b, see Fig. 2B). In each trial, a word, either “RED” or “BLUE,” indicating the color of the items to be attended to (the selection instruction), was presented for 200 ms, followed by a 100-ms interval. Subsequently, 10 bars were displayed for 250 ms, with half of them in the designated color, creating a set size 5 condition. After a 900-ms delay, only the bars in the attended color reappeared. During the test phase, one of the bars changed its orientation compared to the encoding phase. Participants were required to identify which one of the five bars had changed its orientation. This Filtering Localization phase comprised a total of 60 trials. Our behavioral measure of interest was the accuracy in the Filtering Change Localization task.

##### Attention Control Tasks

###### (1) Flanker Square

The Flanker Square task was adapted from the Flanker task (Burgoyne et al., 2023, see Fig. 2C). In each trial, participants were presented with a target stimulus and two possible response options, both consisting of sets of five arrows arranged horizontally (e.g., <<><<). Participants were instructed to choose the response option where the middle arrow aligned in direction with the outer arrows of the target stimulus. For example, if the target stimulus displayed arrows pointing left and right (e.g., <<><<), participants were to select the response option with a central arrow pointing left (e.g., >><>>). Thus, the task required participants to focus on the outer arrows of the target stimulus and the central arrow of the response options, while ignoring the central arrow of the target stimulus and the outer arrows of the response options. Each participant completed 30 seconds of practice followed by a 90-second test phase. The Flanker Square score was calculated as the difference between the number of correct and incorrect responses. Higher scores in Flanker Square corresponded to better attentional control abilities.

###### (2) Simon Square

The Simon Square task was adapted from the Simon task (Burgoyne et al., 2023, see Fig. 2D). In each trial, participants were presented with a target stimulus, an arrow, and two response options: the words “RIGHT” and “LEFT.” Participants were instructed to select the response option that matched the direction indicated by the arrow. For example, if the arrow pointed to the left, participants would choose the response option with the word “LEFT.” The target stimulus and response options could appear on either side of the screen with equal probability. Therefore, participants needed to focus on the direction of the arrow while interpreting the response options’ meaning, ignoring the screen’s side where the stimuli appeared. Each participant completed 30 seconds of practice followed by a 90-second test phase. The Simon Square score was calculated as the difference between the number of correct and incorrect responses. Higher scores in Simon Square corresponded to better attentional control abilities.

### Preprocessing and artifact rejection of EEG signals

Participants were positioned in an electrically isolated booth, with their heads stabilized using a cushioned chinrest placed 74 cm from the display screen. Electroencephalographic (EEG) signals were recorded via 30 active silver/silver chloride (Ag/AgCl) electrodes integrated into a stretchable cap (actiCHamp system, Brain Products, Munich, Germany), arranged according to the international 10-20 placement standard (electrode sites: Fp1, Fp2, F7, F8, F3, F4, Fz, FC5, FC6, FC1, FC2, C3, C4, Cz, CP5, CP6, CP1, CP2, P7, P8, P3, P4, Pz, PO7, PO8, PO3, PO4, O1, O2, Oz). Additional electrodes were adhered to the left and right mastoids using adhesive stickers, and a ground electrode was integrated at the Fpz site within the cap. EEG signals were initially referenced to the right mastoid and subsequently re-referenced offline to the average of both mastoids. The signal was bandpass filtered (0.01–80 Hz) with a 12 dB/octave roll-off and digitized at a sampling rate of 500 Hz. All electrode impedances were maintained below 10 kΩ.

To track ocular activity such as blinks and saccades, both electrooculographic (EOG) and eye-tracking data were recorded. EOG was captured using five passive Ag/AgCl electrodes: two for vertical EOG (above and below the right eye), two for horizontal EOG (approximately 1 cm lateral to each eye), and one ground electrode on the left cheek. Eye position was also monitored with a desktop-mounted EyeLink 1000 Plus system (SR Research, Ontario, Canada), operating at a sampling rate of 1,000 Hz.

To enhance spatial specificity and reduce volume conduction effects, we applied the surface Laplacian (current source density, CSD) transformation to the EEG data using the spherical spline method. This approach estimates the second spatial derivative of the scalp potential, effectively acting as a spatial high-pass filter and improving signal-to-noise ratio (SNR) for inter-electrode correlation analyses. CSD was computed using the compute_current_source_density function from MNE-Python (Cohen, 2014; Pernier et al., 1988; Perrin et al., 1989) based on the 3D locations of 28 scalp electrodes.

### Artifact Rejection

For horizontal eye movements, we employed a sliding-window algorithm to identify horizontal eye movements using both horizontal electrooculogram (HEOG) signals and eye-tracking gaze data. For the HEOG-based detection, a split-half sliding-window method was applied with a 100 ms window moving in 10 ms increments. An eye movement was flagged if the voltage difference between the two halves of the window exceeded 20 µV. This HEOG-based artifact detection was used only in trials where eye-tracking data were unreliable or unavailable. In parallel, eye-tracking rejection was carried out by analyzing horizontal (x-axis) and vertical (y-axis) gaze positions using the same 100 ms window and 10 ms step size. A trial was excluded if eye position shifted more than 0.5° of visual angle within any given window.

To detect blinks, we applied a sliding-window analysis to the vertical EOG signal using an 80 ms window and a 10 ms step size. A blink was marked when the voltage change across the window exceeded 30 µV. Complementarily, we identified blink periods in the eye-tracking data by locating segments where positional data were missing, indicating that the eyes were closed.

### Inter-electrode EEG Correlations

We measure correlations between the mean trial time courses between all pairs of electrodes (Hakim et al., 2021). To do so, we averaged the raw amplitude at each of our non-reference 28 electrodes across all trials from time points 0 (the onset of the memory array) to 800 ms, which is the longest artifact-free time for each trial during the coding phase. Since we have four set size 32 blocks and one set size 128 block, the number of trials during the encoding phase was the same for both conditions, so the signal-noise-ratio of the resulting event-related potential (ERP) was similar between our two different conditions. We next computed the Pearson correlation of this trial-averaged ERP for all pairwise electrodes for each participant separately. For each participant, this resulted in a 28 x 28 matrix of the correlation between the time course of each electrode to each other electrode.

### EEG Fingerprinting

The fingerprinting analysis examined whether individuals’ inter-electrode correlation patterns were both unique and stable enough to distinguish them from others in the group and thus could potentially serve as a trait-level neural signature of long-term memory performance. This approach was adapted from work on functional connectome fingerprinting using fMRI (Finn et al., 2015). To identify individuals in our dataset, we compared each participant’s vectorized inter-electrode correlation matrices from their set size 32 and set size 128 conditions. If these patterns are unique between and stable within individuals, we would expect a participant’s set size 32 and set size 128 correlation patterns to be more similar to each other than to those of other participants. To test this, we correlated each participant’s set size 32 vector of inter-electrode correlations (the “target”) with a database of all participants’ set size 128 vectors. An individual was considered correctly identified if their highest correlation was with their own set size 128 data. We repeated the analysis with the roles reversed, using set size 128 vectors as targets and set size 32 vectors as the database. Fingerprinting accuracy was then calculated as the proportion of correctly identified individuals out of the total number of participants.

### EEG-based Predictive Modeling

Prediction methods were adopted from previous work on connectome-based predictive modeling (Finn et al., 2015, p. 201; Rosenberg et al., 2016) as described in Hakim et al. (2021). We first separated data into training and testing sets. For all of our models, we used 10-fold cross-validation and permutation testing to determine whether the model significantly predicted behavior.

To identify the features that were significantly related to behavior (for instance, hit rate) in the training sample, we correlated each value in the inter-electrode correlation matrix to behavior using Spearman’s correlation. Due to the sparse nature of the EEG interelectrode connectivities (Hakim et al., 2021), we used a threshold of p = 0.2 that most strongly predicted behavior in positive and negative directions. This approach allows us to compare the inter-electrode correlation feature sets (‘‘networks’’) that positively and negatively predict behavior.

Using these selected features, we then calculated single-subject summary values for all training subjects. To do this, we summed the correlation-strength values for each individual for both the positively and negatively predictive feature sets separately, and then we took the difference between them. We trained a linear model to predict behavior performance from these summary features in the training set. We then used this model to predict behavior performance (i.e., hit rate) in the testing set. To do this, we calculated the summary feature in each test set participant and input these summary scores into the model defined in the training sample to generate predictions for each test set subject’s behavior score.

To determine whether our models significantly predicted performance, we compared predicted and observed behavior scores using Spearman rank correlation (rho) values and non-parametric p-values the correlations with 5,000 times of iterations. This assesses whether our model successfully predicts which participants have relatively better or worse encoding performance. However, model significance remains consistent when predictions are evaluated with Pearson correlation. We additionally calculated null rho_null values within each fold by shuffling the behavioral labels. This resulted in 5,000 observed and 5,000 null rho values. We took the mean of the z-transformed rho values, resulting in one observed rho value across all folds and iterations. To determine whether the observed rho value was significantly above change, we compared the rho_null distribution to mean observed rho using the following formula:

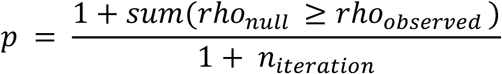

where rho_null is the null rho distribution; rho_observed is the mean rho value from the observed sample; n_interation is the number of iterations; and p is the non-parametric p-value for rho.

## Results

Our primary objective was to determine whether EEG inter-electrode correlations during long-term memory encoding predict individual differences in behavioral performance. Prior to examining brain–behavior relationships, we first assessed recognition accuracy (i.e., hit rates) for old images in the set size 32 and set size 128 conditions. As shown in **Figure 3A**, there were substantial individual differences in performance across both conditions, and hit rates were strongly correlated between the two set sizes (r(109) = 0.80, p < 0.01), suggesting that both tasks tap into a shared underlying long-term memory ability.

**Fig 3.**
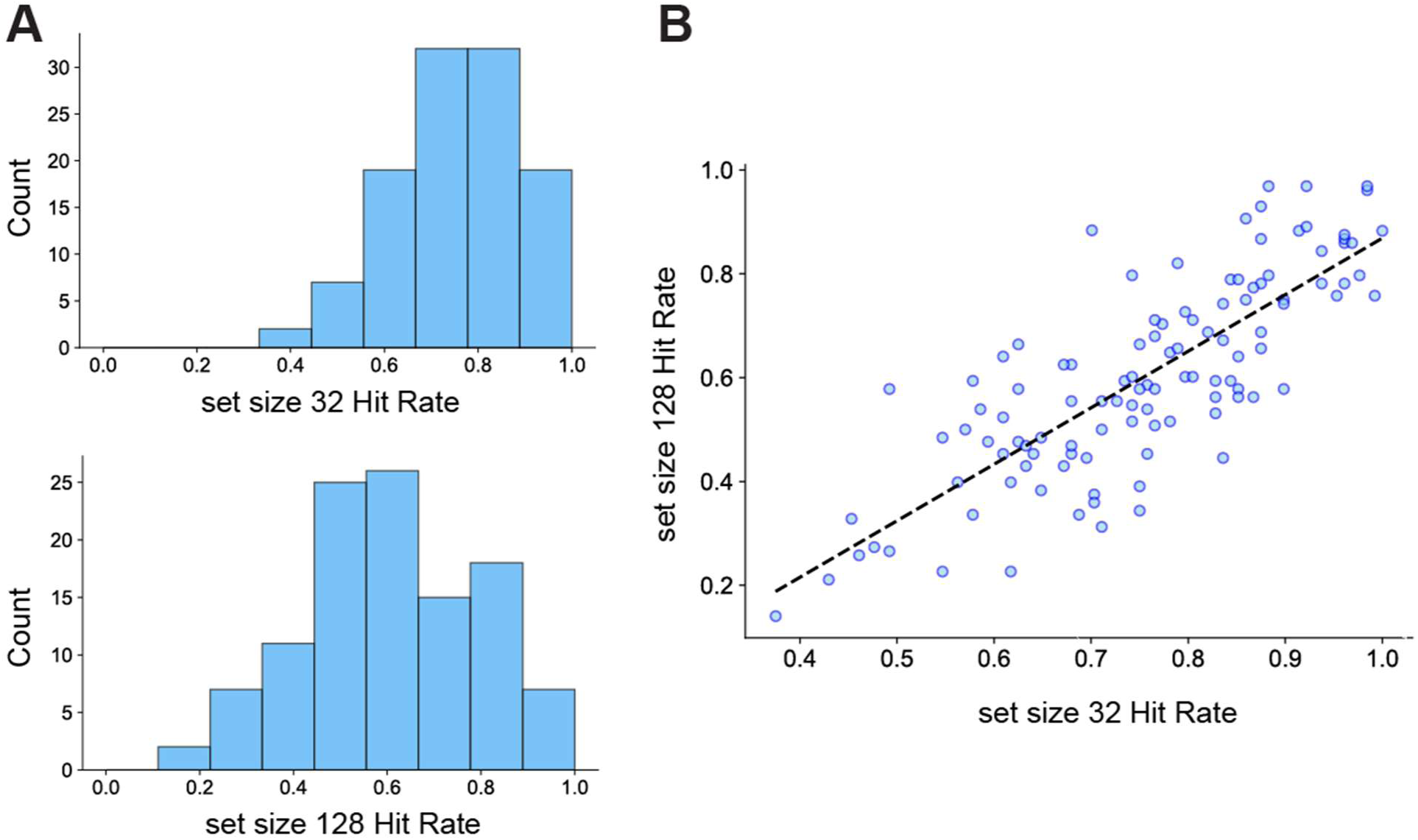
Visual Long-Term Memory Behavior Results. (A) Distribution of hit rates for set size 32 and 128. (B) Individual differences in LTM abilities, measured by hit rates, were highly correlated between the two set sizes.

To validate the stability and subject-specificity of the neural measures, we tested whether inter-electrode correlation patterns identify individuals across the two set-size conditions. For each participant, we computed inter-electrode correlation matrices separately for set size 32 and set size 128 trials. An EEG fingerprinting approach (Hakim et al., 2021) successfully identified 102 out of 111 participants (91.89%) when set size 32 was used as the template and set size 128 as the target, and 108 out of 111 participants (97.30%) when the template and target were reversed. These identification rates, far above chance level (1/111), indicate that inter-electrode correlations are highly reliable and individually distinctive.

As a control analysis, we evaluated whether simple amplitude-based measures achieved similar identification accuracy. Using the mean amplitude at each electrode, fingerprinting performance dropped significantly, identifying only 53.15% and 50.45% of participants when using set size 32 and set size 128 as templates, respectively. These results highlight that inter-electrode correlation patterns uniquely capture individual-specific neural signatures, making them particularly well-suited for individual differences analyses in the context of visual long-term memory encoding.

### Interelectrode Correlation with Encoding-Phase EEG predicts Visual LTM Differences

With our fingerprinting analysis, we confirmed that interelectrode correlations during a LTM encoding phase are sensitive and reliable measures of participant uniqueness. We next investigated whether these correlations could predict behavioral differences in memory performance. We employed connectome-based predictive modeling (CPM) using a 10-fold cross-validation procedure, selecting the interelectrode correlations correlated with LTM encoding performance at p < 0.2. Using 5,000 permutations to generate a null distribution, we found that interelectrode correlations during the encoding phase significantly predicted individual differences in hit rates for both the set size 32 (r(109) = 0.29, p < 0.01, **Fig. 4**) and set size 128 (r(109) = 0.30, p < 0.01, **Fig. 4**) conditions.

**Fig 4.**
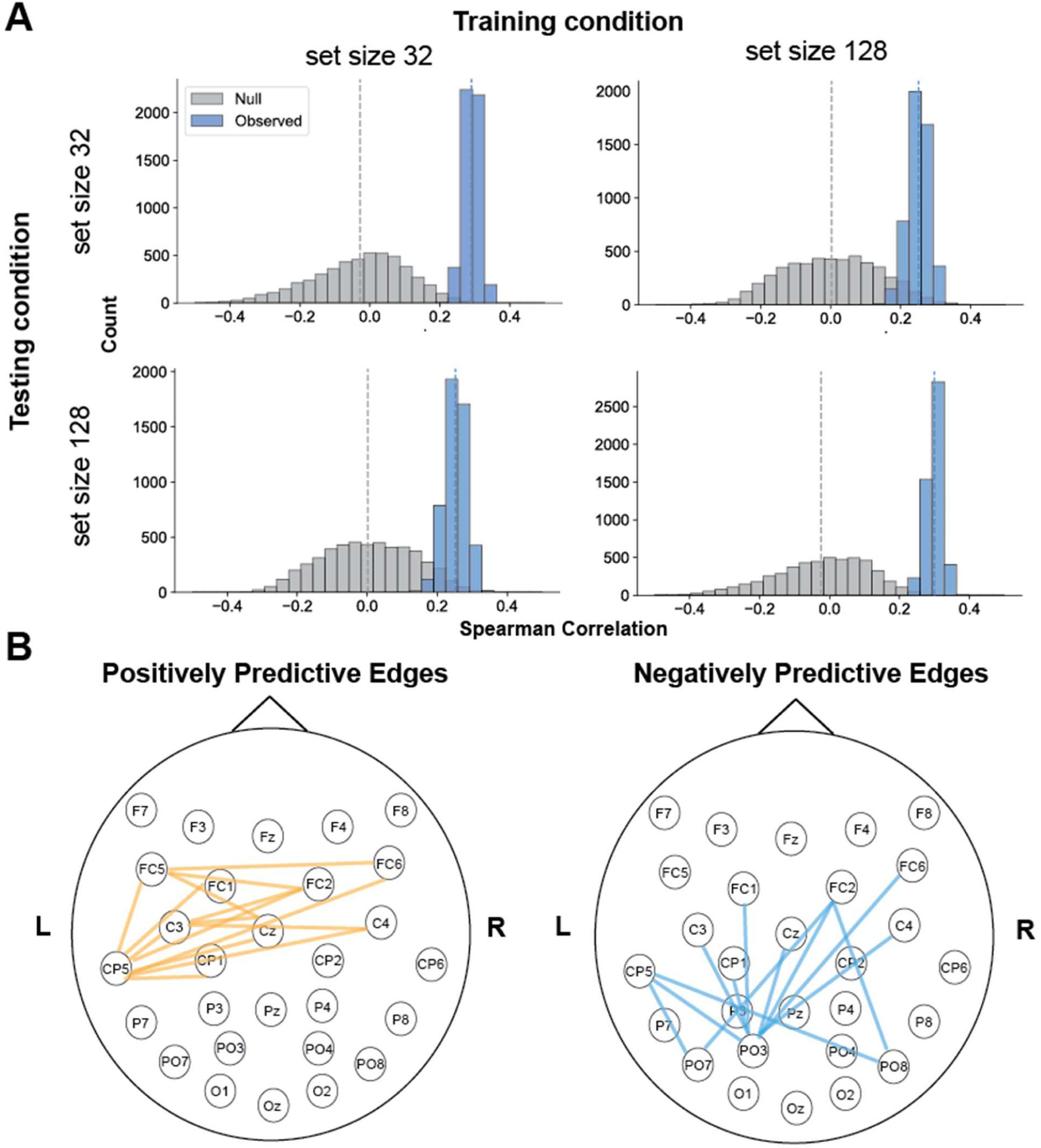
EEG CPM predicted individual differences in LTM encoding with both set sizes. (A) EEG interelectrode correlation model significantly predicts individual differences in LTM encoding abilities for both set size 32, set size 128, as well as cross training between the two set sizes. (B) Significant edges overlapping between set size 32 and set size 128 across EEG electrodes predicting LTM differences (orange represents positively predictive edges, while blue represents negatively predictive edges).

To assess whether the model captured a generalizable LTM encoding ability rather than condition-specific variance, we conducted cross-condition prediction analyses. Specifically, we trained the model on EEG data from the set size 32 condition and tested it on left-out participants’ EEG data from the set size 128 condition. Predictions remained significant, indicating successful generalization (r(109) = 0.25, p = 0.02, **Fig. 4**). The reverse analysis, training on set size 128 and testing on set size 32, also yielded significant predictions (r(109) = 0.25, p = 0.02, **Fig. 4**). These cross-condition results suggest that the inter-electrode correlation model captures core individual differences in LTM encoding, independent of set size (cross-training rho values not different from within-training rho values, nonparametric *p*s > 0.83).

We showed that we can successfully predict individual differences in long-term memory encoding using interelectrode correlation from a single condition (either set size 32 or 128). A remaining question is whether averaging more trials, across both set sizes 32 and 128, will better maximize our theoretical predictive power to detect these individual differences, given that prior ERP research emphasizes the importance of trial count for reliable analysis (Boudewyn et al., 2018). To examine this possibility, we combined data from both set sizes by averaging the time courses across all encoding trials. Inter-electrode correlations were then recomputed from this combined dataset, and the resulting model significantly predicted the average hit rate across both conditions (r(109) = 0.31, p < 0.01). Together, these results demonstrate that EEG-based inter-electrode correlation patterns during encoding robustly capture individual differences in long-term memory performance across 111 participants (**Fig. 5B**, interelectrode correlation patterns for LTM encoding).

**Fig 5.**
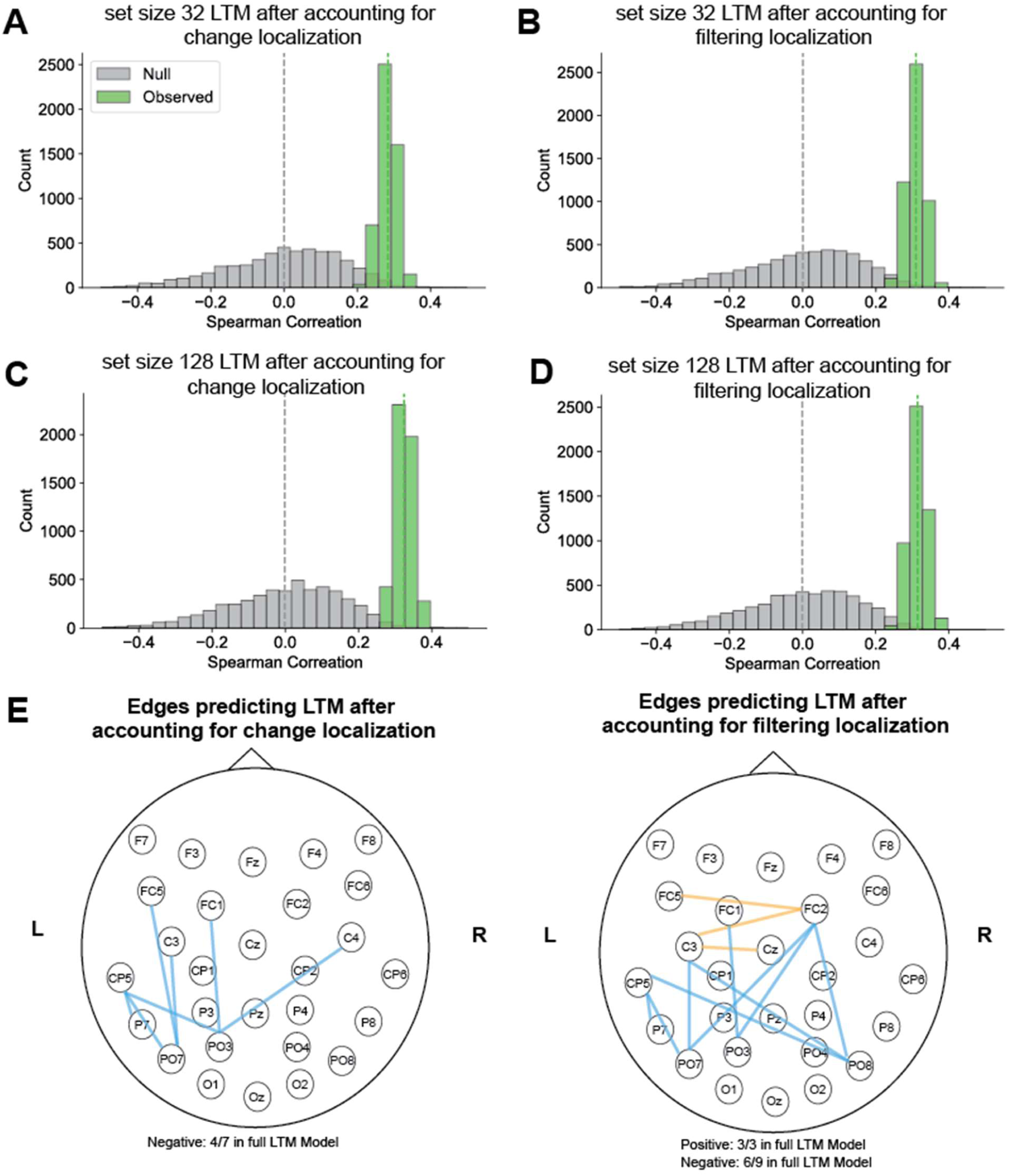
EEG CPM predicted individual differences in LTM encoding after regressing out Visual Working Memory differences. (A-D) EEG interelectrode correlation model significantly predicts individual differences in LTM encoding with working memory (VWM) tasks regressed out (orange represents positively predictive edges, while blue represents negatively predictive edges). (E-F) Significant edges across EEG electrodes predicting LTM differences when VWM differences were regressed out.

### Encoding-Phase EEG Synchrony Predicts Individual Differences in Visual Long-Term Memory After Controlling for Visual Working Memory

We next tested if these neural signatures predict LTM encoding independent of visual working memory (VWM) capacity. We analyzed data from a subset of 80 participants who completed both the EEG and online behavioral sessions. Behaviorally, we found that LTM performance in the set size 32 condition significantly predicted VWM ability, as measured by change localization (r(78) = 0.37, p < 0.01) and filtering localization (r(78) = 0.27, p = 0.01). Similarly, LTM performance in the set size 128 condition significantly predicted change localization accuracy (r(78) = 0.25, p = 0.02), and marginally predicted filtering localization (r(78) = 0.21, p = 0.05).

To determine whether our LTM model predicted LTM-specific variance rather than variance shared with VWM, we conducted a regression-based control analysis. For the training set in each cross-validation fold, we regressed VWM scores from LTM performance and extracted the residuals, creating an LTM measure that was independent of VWM ability. We then applied the regression function trained with residuals of LTM after regressing out VWM differences to the test set, generating predicted residuals of LTM performance. This procedure ensured the independence between training and test data, such that the regression function parameters were not affected by any of the hold-out test data. If the EEG-based CPM captures LTM-specific variance, it should still predict these residualized scores. Supporting this specificity, the model trained on residualized LTM performance significantly predicted residual LTM scores in the test participants for both set size 32 (regress out change localization: r(78) = 0.28, p = 0.01, **Fig. 5A**; regress out filtering localization: r(78) = 0.33, p < 0.01, **Fig. 5B**) and set size 128 (regress out change localization: r(78) = 0.31, p < 0.01, **Fig. 5C**; regress out filtering localization: r(78) = 0.31, p < 0.01, **Fig. 5D**) conditions. These findings demonstrate that EEG inter-electrode correlations during encoding capture variance that is specific to long-term memory ability, above and beyond shared variance with visual working memory.

### Encoding-Phase EEG Synchrony Predicts Individual Differences in Visual Long-Term Memory After Controlling for Attentional Control

Other than WM abilities, another potentially relevant construct is attentional control (AC, Martin et al., 2021), as prior research has shown moderate correlations between AC and LTM performance (Zhao & Vogel, 2025b). To explore this possibility, we analyzed data from a subset of 80 participants who completed both the EEG and online behavioral sessions. LTM performance in the set size 32 condition significantly predicted AC ability, as measured by accuracies in the Flanker Square task (r(78) = 0.23, p = 0.03) and accuracies in the Simon Square task (r(78) = 0.21, p = 0.05). Similarly, LTM performance in the set size 128 condition was positively, although not statistically significantly, correlated to Flanker Square accuracy (r(78) = 0.13, p = 0.24), and Simon Square accuracy (r(78) = 0.18, p = 0.10).

Replicating our findings with VWM, the model trained on residualized LTM performance significantly predicted residual LTM scores in the test participants for both set size 32 (regress out Flanker Square: r(78) = 0.31, p < 0.01, **Fig. 6A**; regress out Simon Square: r(78) = 0.34, p < 0.01, **Fig. 6B**) and set size 128 (regress out Flanker Square: r(78) = 0.33, p < 0.01, **Fig. 6C**; regress out Simon Square: r(78) = 0.34, p < 0.01, **Fig. 6D**) conditions.

**Fig 6.**
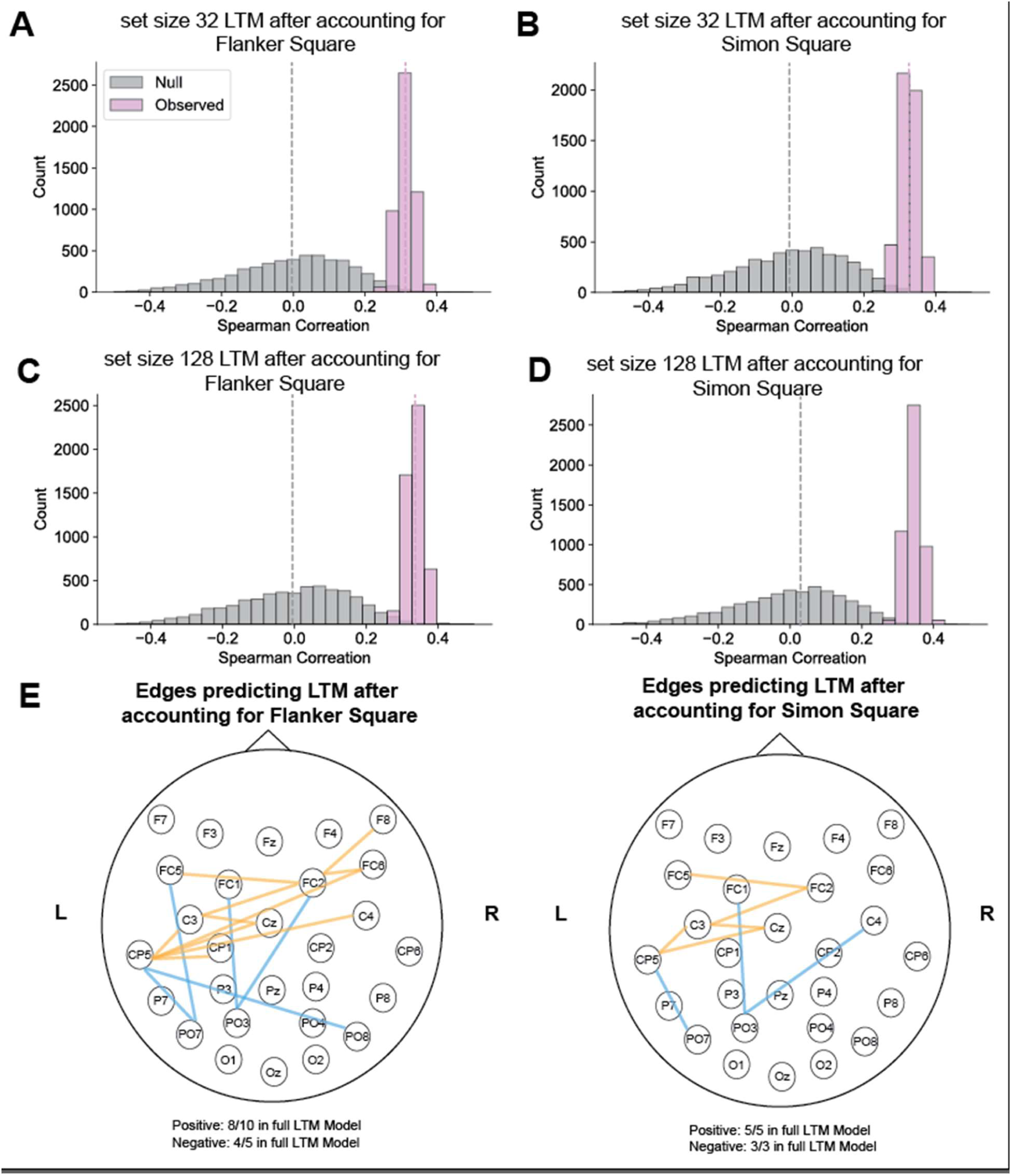
EEG CPM predicted individual differences in LTM encoding after regressing out Attentional Control differences. (A-D) EEG interelectrode correlation model significantly predicts individual differences in LTM encoding with attentional control (AC) tasks regressed out. (E-F) Significant edges across EEG electrodes predicting LTM differences when AC differences were regressed out (orange represents positively predictive edges, while blue represents negatively predictive edges).

### Spatial Distribution of Edges in LTM Encoding-Phase Interelectrode Correlation Models

In the full LTM encoding model (**Figure 4B**), we observed that edges positively and negatively predicting performance were largely in left parietal electrodes. Specifically, the positively predictive correlations covered the right parietal frontal electrodes, and negatively predictive correlations covered more posterior parietal to occipital electrodes. These observations suggest that left parietal electrodes may be a key hub in predicting individual differences in LTM encoding abilities.

The interelectrode correlation patterns for the residual LTM encoding models, after regressing out VWM capacity, were sparser than the full model, but the remaining significant correlations highly overlapped with the full model (regressing out change localization: 57.14% in the full model, 100% within 2.5 cm/1 electrode in the full model; regressing out filtering localization: 75.00% in the full model, 100% within 2.5 cm/1 electrode of the full model, **Figure 5E**). We observed similar patterns with AC abilities (regressing out Flanker Square: 80.00% in the full model, 100% within 2.5 cm/1 electrode in the full model; regressing out Simon Square: 100.00% in the full model, **Figure 6E**). Thus, interelectrode correlations with left parietal electrodes remain uniquely predictive of LTM encoding even regressing out VWM and AC abilities.

### Encoding-Phase EEG Interelectrode Correlation Predicts Individual Differences in Visual Long-Term Memory throughout the Encoding Phase

We next sought to determine when during the encoding period the model began to significantly predict individual differences in visual long-term memory (LTM). Around or before 400 ms post-stimulus, prior ERP studies have shown that subsequently remembered items are associated with a frontal positivity. In contrast, after 500 ms, the subsequent memory effect tends to manifest as a posterior-going positivity (e.g., N400; Kutas & Hillyard, 1980; Mecklinger & Kamp, 2023; Meng et al., 2014; Sundby et al., 2019). However, these studies did not examine whether these neural signatures predict individual differences in long-term memory (LTM) encoding. Therefore, a time-resolved analysis could address two key questions: (1) Do early and late neural signals contribute to individual differences in LTM encoding? and (2) If both early and late signals are present, do they share similar inter-electrode correlation patterns (i.e., similar edges)?

To address this, we first performed a temporal analysis by segmenting each trial into 100-ms windows, using a 25-ms jumping window to compute inter-electrode correlation matrices across time. For each time point, we conducted a 5,000-iteration permutation test, with window start times spanning from 175 ms before stimulus onset to 700 ms post-stimulus onset. At each window, CPM models were trained and tested using the inter-electrode correlations derived from the 100-ms segment centered on that specific time point, as illustrated in **Fig. 7A**.

**Fig 7.**
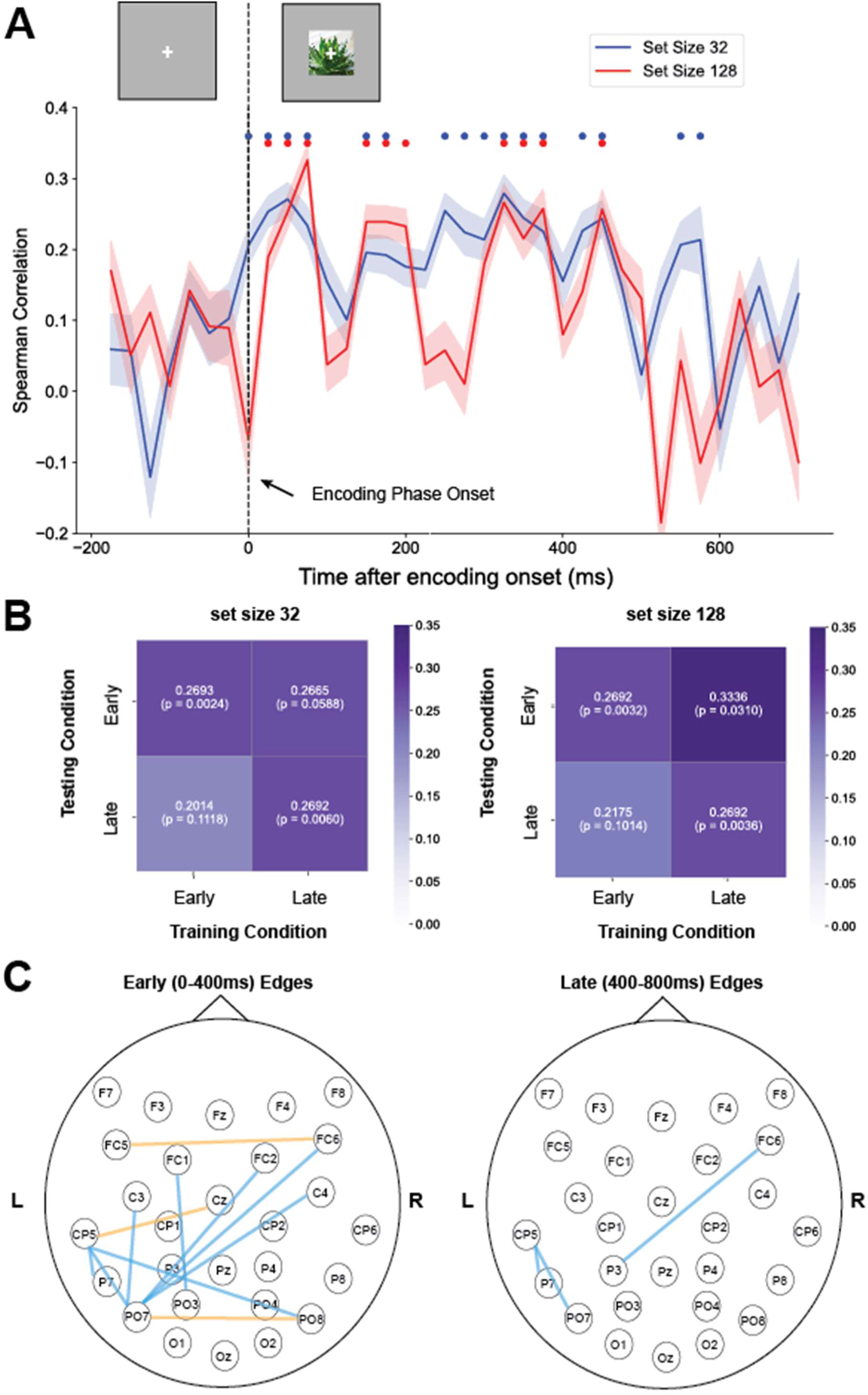
EEG CPM predicted individual differences in LTM encoding across a wide temporal window. (A) EEG interelectrode correlation patterns predict LTM differences across time (correlation between predicted LTM and observed LTM). EEG encoding phase data was divided into 100 ms epochs, with 25-ms steps, from 200 ms before the stimulus onset to 800 ms after stimulus onset. The predictive power with set size 32 and 128 both extended from 0 ms to 500-600 ms after the stimulus onset. The significance dots reflect uncorrected p < 0.05. (B) Cross training (early: 0-400 ms versus late: 400-800ms) correlations of LTM differences with set size 32 and 128 data. The diagonal showed within-condition training and testing, versus the off-diagonal columns reflected cross-condition training and test (i.e., train on early and test on late window and vice versa). (C) Significant edges predicting LTM differences trained with early versus late window EEG encoding data (orange represents positively predictive edges, while blue represents negatively predictive edges).

We first confirmed that no significant brain–behavior correlations were observed prior to stimulus onset for either set size condition, indicating that the model’s predictive performance was not driven by pre-trial neural synchrony between electrode ERPs. Shortly after stimulus onset, the models began to reliably predict individual differences in LTM. For set size 32, significant prediction first emerged in the 0–100 ms window, while for set size 128, it emerged slightly later, in the 25–125 ms window. Notably, significant prediction persisted throughout much of the encoding period, with the final time windows showing significance between 575–675 ms for set size 32 and 450–550 ms for set size 128 (see **Fig. 7A**).

After establishing that both early and late time windows in the trial show inter-electrode correlations that predict LTM encoding, we next asked whether these predictions rely on similar neural patterns. To test this, we performed a cross-training analysis between early (0-400 ms) and late (400-800 ms) trial segments. If both time windows rely on shared neural information, then training a model on one segment should allow accurate prediction from the other. The results of the cross-training analysis revealed an asymmetry: when we trained on the late trial segment and tested on the early segment, the model was largely successful in predicting individual differences in LTM encoding (set size 32: r(109) = 0.27, p = 0.06; set size 128: r(109) = 0.33, p = 0.03). However, training on the early segment and testing on the late segment failed to produce accurate predictions (set size 32: r(109) = 0.20, p = 0.11; set size 128: r(109) = 0.22, p = 0.10, not significantly different from train later test early, *p*s > 0.60, see **Fig. 7B**). To better understand this asymmetry, we examined the positive and negative inter-electrode correlation edges for early and late trials (**Fig. 7C**). We found that the early trials exhibited a greater number of both positive and negative edges, whereas the late trials had only three significant negative edges. Importantly, two of these three late-trial edges were identical to those seen in the early-trial models, and the third was spatially adjacent with the other (P3 versus PO7, ∼2.5 cm from each other). These findings suggest that the early phase (0-400 ms) of the trial is characterized by richer and more distributed inter-electrode correlations that are predictive of LTM encoding differences. In contrast, the late phase (400-800 ms) appears to contribute more selectively, primarily through a small set of negative correlations.

These findings suggest that the predictive power of the CPM is tied specifically to stimulus encoding, rather than baseline pre-encoding activity, and that individual differences in VLTM are encoded in neural activity sustained throughout the first several hundred milliseconds following stimulus presentation.

## Discussion

In our study, we investigated whether interelectrode correlations predict individual differences in long-term memory encoding, using set sizes of 32 and 128, both well beyond typical working memory capacity (3-4 items). We demonstrated that interelectrode correlation patterns could identify individuals with a fingerprinting-style analysis, achieving over 90% accuracy. Furthermore, patterns of interelectrode correlations during LTM encoding predicted individual differences in performance on both the set size 32 and 128 tasks. Cross-training between the two set sizes generalized effectively, indicating that the neural patterns reflected a more generalized LTM encoding ability rather than performance on a single recognition task. To assess whether this network was specific to LTM encoding, we further examined whether interelectrode correlations could predict LTM performance even after accounting for other cognitive abilities such as visual working memory (VWM) and attentional control (AC). Using residual LTM performance, after regressing out scores from four separate WM and AC tasks, we found that our model still significantly predicted LTM encoding. This suggests that we identified a neural signature specifically associated with LTM encoding, independent of other cognitive domains.

Looking into the spatial distribution of edges significantly predicting individual differences in LTM encoding, both positively and negatively predictive interelectrode correlations covered left parietal electrodes. Whereas positively predictive edges were distributed more towards anterior frontal to parietal electrodes, negatively predictive edges were shifted to more posterior parietal and occipital electrodes. The residual models, built with LTM encoding abilities with VWM or AC regressed out, contained a subset of edges of the full LTM encoding model, suggesting that these models were successful in uniquely targeting a sparser network that predicts LTM encoding differences but not VWM or AC abilities. The residual models retained the negative edges from left parietal electrodes and some positive centro-parietal edges. Interestingly, the parietal electrodes here are involved in the ERP subsequent memory effect, in that a more positive waveform was observed during encoding when items were memorized compared to when they were later forgotten (Folgueira-Ares et al., 2017; Guo et al., 2005; Mangels et al., 2010; Paller et al., 1987; Sanquist et al., 1980; Yovel & Paller, 2004). Moreover, the parietal-frontal interelectrode correlations observed in our study also align with the ERP literature that two subsequent memory effects emerged within 300 - 600 ms after stimulus onset, one peaked at frontal electrodes and the other peaked at the parietal electrodes (i.e., Otten & Rugg, 2001; see review, Friedman & Johnson Jr, 2000). Altogether, our findings showed a relatively sparse set of edges that specifically and efficiently captured individual differences in LTM encoding.

Finally, we explored the temporal dynamics of these neural processes. Interelectrode correlation patterns did not predict activity prior to trial onset, indicating that predictive information emerged with memory encoding rather than from averaged pre-stimulus state for each individual. Although non-trial-locked signals such as pretrial attentional states (Adam et al., 2015; Corriveau et al., 2025) may still affect performance in individual trials, we cannot detect them here because we employed averaged ERP interelectrode correlations to enhance the signal-to-noise ratio. Future research is needed to evaluate the effectiveness of single-trial EEG interelectrode correlations in predicting cognitive performance. In the current study, predictive power persisted over a broad time window of approximately 500-600 ms post-stimulus onset. The differences between early and late EEG interelectrode correlation patterns suggest that dynamic processes unfolding over the course of stimulus presentation affect the encoding of items into LTM, aligning with within-subject findings from event-related potential (ERP) studies (e.g., N400; Kutas & Hillyard, 1980; Mecklinger & Kamp, 2023; Meng et al., 2014; Sundby et al., 2019). A further look into the predictive model features showed that, early in the trial, predictive interelectrode correlations spanned parietal and frontal electrodes, potentially reflecting the engagement of early sensory-related encoding processes at the onset of memory stimuli. In contrast, later predictive features were sparser and predominantly negatively predictive of long-term memory. These correlations included parietal and parieto-frontal electrodes overlapping with those implicated in the late parietal effect observed in the subsequent memory literature (see review, Friedman & Johnson Jr, 2000; Mecklinger & Kamp, 2023).

One possible cognitive factor underlying the observed neural dynamics in our LTM model is that, as list lengths exceeded VWM capacity, participants may have begun to shift their organizational strategies (Stoff & Eagle, 1971). Interestingly, such dynamic strategy shifts have been shown to enhance LTM performance, yet they are not strongly tied to individual differences in VWM capacity (Unsworth, 2016). In other words, participants’ decisions to flexibly adjust their strategies could explain why our neural model uniquely captures LTM encoding differences.

More broadly, our study supports the idea that long-term memory encoding involves processes distinct from those used in working memory encoding. The specificity of our model’s predictions for LTM, after controlling for VWM and attentional control, suggests the presence of an LTM-specific factor that operates independently of VWM and AC abilities. This aligns with findings from prior behavioral research (Robison et al., 2024; Unsworth, 2019; Zhao et al., 2025; Zhao & Vogel, 2025b, 2025a) and fMRI studies (Lin et al., 2021; Zhou et al., 2025). Future behavioral and neural studies are needed to better understand the distinct mechanisms underlying LTM encoding. A combined approach that integrates both correlational and experimental methods, as recommended by Unsworth (2019), may be especially useful in advancing our understanding of individual differences in LTM.

## Author Contributions

Chong Zhao played a lead role in conceptualization, data curation, formal analysis, investigation, methodology, writing–original draft, and writing– review and editing. Edward K. Vogel played a lead role in conceptualization, funding acquisition, resources, supervision, and writing–review and editing. Monica D. Rosenberg played a lead role in conceptualization, formal analysis, investigation, methodology, funding acquisition, resources, supervision, and writing–review and editing.

## Reference

Adam, K. C., Mance, I., Fukuda, K., & Vogel, E. K. (2015). The contribution of attentional lapses to individual differences in visual working memory capacity. Journal of Cognitive Neuroscience, 27(8), 1601–1616.

Atkinson, R. C., & Shiffrin, R. M. (1968). Human Memory: A Proposed System and its Control Processes. Psychology of Learning and Motivation - Advances in Research and Theory, 2(C), 89–195. 10.1016/S0079-7421(08)60422-3

Boudewyn, M. A., Luck, S. J., Farrens, J. L., & Kappenman, E. S. (2018). How many trials does it take to get a significant ERP effect? It depends. Psychophysiology, 55(6), e13049.

Brady, T. F., Konkle, T., Alvarez, G. A., & Oliva, A. (2008). Visual long-term memory has a massive storage capacity for object details. Proceedings of the National Academy of Sciences of the United States of America, 105(38), 14325–14329. 10.1073/pnas.0803390105

Burgoyne, A. P., Tsukahara, J. S., Mashburn, C. A., Pak, R., & Engle, R. W. (2023). Nature and measurement of attention control. Journal of Experimental Psychology: General.

Cohen, M. X. (2014). Analyzing neural time series data: Theory and practice. MIT press.

Corriveau, A., Rosenberg, M., deBettencourt, M., & deBettencourt, M. T. (2025). Cognitive Neuroscience of Attention and Memory Dynamics.

Cowan, N. (2001). The magical number 4 in short-term memory: A reconsideration of mental storage capacity. Behavioral and Brain Sciences, 24(1), 87–114. 10.1017/S0140525X01003922

Finn, E. S., Shen, X., Scheinost, D., Rosenberg, M. D., Huang, J., Chun, M. M., Papademetris, X., & Constable, R. T. (2015). Functional connectome fingerprinting: Identifying individuals using patterns of brain connectivity. Nature Neuroscience, 18(11), 1664–1671.

Folgueira-Ares, R., Cadaveira, F., Rodríguez Holguín, S., López-Caneda, E., Crego, A., & Pazo-Álvarez, P. (2017). Electrophysiological anomalies in face–name memory encoding in young binge drinkers. Frontiers in Psychiatry, 8, 216.

Friedman, D., & Johnson Jr, R. (2000). Event-related potential (ERP) studies of memory encoding and retrieval: A selective review. Microscopy Research and Technique, 51(1), 6–28.

Fukuda, K., & Vogel, E. K. (2019). Visual short-term memory capacity predicts the “bandwidth” of visual long-term memory encoding. Memory and Cognition, 47(8), 1481–1497. 10.3758/s13421-019-00954-0

Fukuda, K., & Woodman, G. F. (2015). Predicting and Improving Recognition Memory Using Multiple Electrophysiological Signals in Real Time. Psychological Science, 26(7), 1026–1037. 10.1177/0956797615578122

Guo, C., Voss, J. L., & Paller, K. A. (2005). Electrophysiological correlates of forming memories for faces, names, and face–name associations. Cognitive Brain Research, 22(2), 153–164.

Hakim, N., Awh, E., Vogel, E. K., & Rosenberg, M. D. (2021). Inter-electrode correlations measured with EEG predict individual differences in cognitive ability. Current Biology, 31(22), 4998–5008.

Hebart, M. N., Contier, O., Teichmann, L., Rockter, A. H., Zheng, C. Y., Kidder, A., Corriveau, A., Vaziri-Pashkam, M., & Baker, C. I. (2023). THINGS-data, a multimodal collection of large-scale datasets for investigating object representations in human brain and behavior. Elife, 12, e82580.

James, W. (1890). The principles of psychology. Henry Holt.

Kutas, M., & Hillyard, S. A. (1980). Reading senseless sentences: Brain potentials reflect semantic incongruity. Science, 207(4427), 203–205.

Lin, Q., Yoo, K., Shen, X., Constable, T. R., & Chun, M. M. (2021). Functional connectivity during encoding predicts individual differences in long-term memory. Journal of Cognitive Neuroscience, 33(11), 2279–2296.

Lionel, S. (1973). Learning 10 000 pictures. Quarterly Journal of Experimental Psychology, 25(1973), 207–222.

Luck, S. J., & Vogel, E. K. (1997). The capacity of visual working memory for features and conjunctions. Nature, 390(6657), 279–284. 10.1038/36846

Mangels, J. A., Manzi, A., & Summerfield, C. (2010). The first does the work, but the third time’s the charm: The effects of massed repetition on episodic encoding of multimodal face–name associations. Journal of Cognitive Neuroscience, 22(3), 457–473.

Martin, J. D., Tsukahara, J. S., Draheim, C., Shipstead, Z., Mashburn, C. A., Vogel, E. K., & Engle, R. W. (2021). The Visual Arrays Task: Visual Storage Capacity or Attention Control? Journal of Experimental Psychology: General, 150(12), 2525– 2551. 10.1037/xge0001048

Mecklinger, A., & Kamp, S.-M. (2023). Observing memory encoding while it unfolds: Functional interpretation and current debates regarding ERP subsequent memory effects. Neuroscience & Biobehavioral Reviews, 153, 105347.

Meng, Y., Ye, X., & Gonsalves, B. D. (2014). Neural processing of recollection, familiarity and priming at encoding: Evidence from a forced-choice recognition paradigm. Brain Research, 1585, 72–82.

Otten, L. J., & Rugg, M. D. (2001). Electrophysiological correlates of memory encoding are task-dependent. Cognitive Brain Research, 12(1), 11–18.

Paller, K. A., Kutas, M., & Mayes, A. R. (1987). Neural correlates of encoding in an incidental learning paradigm. Electroencephalography and Clinical Neurophysiology, 67(4), 360–371.

Pernier, J., Perrin, F., & Bertrand, O. (1988). Scalp current density fields: Concept and properties. Electroencephalography and Clinical Neurophysiology, 69(4), 385– 389.

Perrin, F., Pernier, J., Bertrand, O., & Echallier, J. F. (1989). Spherical splines for scalp potential and current density mapping. Electroencephalography and Clinical Neurophysiology, 72(2), 184–187.

Robison, M. K., Miller, A. L., Wiemers, E. A., Ellis, D. M., Unsworth, N., Redick, T. S., & Brewer, G. A. (2024). What makes working memory work? A multifaceted account of the predictive power of working memory capacity. Journal of Experimental Psychology: General, 153(9), 2193.

Rosenberg, M. D., Finn, E. S., Scheinost, D., Papademetris, X., Shen, X., Constable, R. T., & Chun, M. M. (2016). A neuromarker of sustained attention from whole-brain functional connectivity. Nature Neuroscience, 19(1), 165–171.

Sanquist, T. F., Rohrbaugh, J. W., Syndulko, K., & Lindsley, D. B. (1980). Electrocortical signs of levels of processing: Perceptual analysis and recognition memory. Psychophysiology, 17(6), 568–576.

Stoff, D. M., & Eagle, M. N. (1971). The relationship among reported strategies, presentation rate, and verbal ability and their effects on free recall learning. Journal of Experimental Psychology, 87(3), 423.

Sundby, C. S., Woodman, G. F., & Fukuda, K. (2019). Electrophysiological and behavioral evidence for attentional up-regulation, but not down-regulation, when encoding pictures into long-term memory. Memory & Cognition, 47(2), 351–364.

Unsworth, N. (2016). The many facets of individual differences in working memory capacity. Psychology of Learning and Motivation, 65, 1–46.

Unsworth, N. (2019). Individual differences in long-term memory. Psychological Bulletin, 145(1), 79.

Unsworth, N., Fukuda, K., Awh, E., & Vogel, E. K. (2014). Working memory and fluid intelligence: Capacity, attention control, and secondary memory retrieval. Cognitive Psychology, 71, 1–26. 10.1016/j.cogpsych.2014.01.003

Vogel, E. K., & Machizawa, M. G. (2004). Neural activity predicts individual differences in visual working memory capacity. Nature, 428(6984), 748–751. 10.1038/nature02447

Wolfe, J. M., Wick, F. A., Mishra, M., DeGutis, J., & Lyu, W. (2023). Spatial and temporal massive memory in humans. Current Biology, 33(2), 405–410.

Yovel, G., & Paller, K. A. (2004). The neural basis of the butcher-on-the-bus phenomenon: When a face seems familiar but is not remembered. Neuroimage, 21(2), 789–800.

Zhao, C., Corriveau, A., Ke, J., Vogel, E. K., & Rosenberg, M. D. (2025). Sustained attention is more closely related to long-term memory than to attentional control. bioRxiv, 2025–03.

Zhao, C., Vogel, E., & Awh, E. (2022). Change localization: A highly reliable and sensitive measure of capacity in visual working memory. Attention, Perception, and Psychophysics. 10.3758/s13414-022-02586-0

Zhao, C., & Vogel, E. K. (2025a). Individual differences in working memory and attentional control continue to predict memory performance despite extensive learning. Journal of Experimental Psychology: General.

Zhao, C., & Vogel, E. K. (2025b). Working memory and attentional control abilities predict individual differences in visual long-term memory tasks. Journal of Memory and Language, 144, 104665.

Zhao, C., & Woodman, G. F. (2020). Converging Evidence That Neural Plasticity Underlies Transcranial Direct-Current Stimulation. Journal of Cognitive Neuroscience, 1–12. 10.1162/jocn_a_01639

Zhou, M. B., Chun, M. M., & Lin, Q. (2025). Modularity Measures of Functional Brain Networks Predict Individual Differences in Long-Term Memory. European Journal of Neuroscience, 61(6), e70052.

